# Diversity analysis and metagenomic insights into the Antibiotic Resistance and Metal Resistances among Hot Spring Bacteriobiome-insinuating inherent environmental baseline levels of Antibiotic and Metal tolerance

**DOI:** 10.1101/773788

**Authors:** Ishfaq Nabi Najar, Mingma Thundu Sherpa, Sayak Das, Nagendra Thakur

## Abstract

Mechanisms of occurrence and expressions of antibiotic resistance genes (ARGs) in thermophilic bacteria are still unknown owing to limited research and data. The evolution and proliferation of ARGs in the thermophilic bacteria is unclear and needs a comprehensive study. In this research, comparative profiling of antibiotic resistance genes and metal tolerance genes among the thermophilic bacteria has been done by culture-independent functional metagenomic methods. Metagenomic analysis showed the dominance of Proteobacteria, Actinobacteria. Firmicutes and Bacteroidetes in these hot springs. ARG analysis through shotgun gene sequencing was found to be negative in case of thermophilic bacteria. However, few of genes were detected but they were showing maximum similarity with mesophilic bacteria. Concurrently, metal resistance genes were also detected in the metagenome sequence of hot springs. Detection of metal resistance gene and absence of ARG’s investigated by whole genome sequencing, in the reference genome sequence of thermophilic *Geobacillus* also conveyed the same message. This evolutionary selection of metal resistance over antibiotic genes may have been necessary to survive in the geological craters which are full of different metals from earth sediments rather than antibiotics. Furthermore, the selection could be environment driven depending on the susceptibility of ARG’s in thermophilic environment as it reduces the chances of horizontal gene transfer. With these findings this article highlights many theories and culminates different scopes to study these aspects in thermophiles.

## Introduction

The generic term “antibiotic” is used to denote any class of chemotherapeutic agents that inhibits or kills microbes (pathogens) by specific interactions with microbial targets [1]. However, the increasing rate of antibiotic resistance in microorganisms is a critical challenge to public health [2–4]. The spread of antibiotic-resistant superbug is posing a threat to morbidity and mortality worldwide [5]. The potential clinical threat of antibiotic-resistant bacteria has abled to gain attention with a focus on the identification and management of environmental reservoirs of antibiotic resistance genes (ARGs). Though antibiotic resistance occurs naturally, human activity including misuse or overuse of antibiotics has exacerbated the scenario. Pathogenic and non-pathogenic bacteria belonging to the mesophilic world has been extensively studied in terms of antibiotic resistance. However, researchers have paid very less attention to antibiotic resistance in thermophilic bacteria. The distribution of ARGs and their antibiotic resistance profile needs attention to avert the possible emergence of an opportunistic pathogen of thermophilic origin. Extensive application of thermophilic bacteria in biotechnological and industrial fields also supports the possibility of future pathogenic emergence with high antibiotic resistance [6]. The most menacing part is that we have no or negligible data and understanding about the mechanisms related to acceptance, occurrence, and expressions of various genes related to antibiotic resistance in thermophilic bacteria. A report on *Legionella* Pneumonia linked to hot spring facility in Gunma prefecture, Japan indicates the same [7].

A wide distribution of heavy metals in nature is also another facet of industrialization and urbanization which has negatively impacted human health [8]. Microorganism has developed metal resistance as a selective evolutionary pressure. Studies widely report the coexistence of ARGs and metal resistance gene (MRGs) in bacteria [9]. Cross-resistance and presence ARGs and MRGs in mobile genetic elements with horizontal transfer efficiency is the underlying mechanism of co-existence. This also makes it important to study the distribution of ARGs alongside MRGs in thermophilic bacteria.

Hot springs are commonly allied with practices like balneotherapy or hydrotherapy. Beliefs regarding the hot springs as sacred entities with various medicinal and healing properties attracted human for a thermal bath [10,11]. These increase the possibility of widespread distribution of ARGs as anthropogenic disturbances in ecosystems play an important role in resistance development. However, infections from hot springs are rare and sporadic. Most of the reported infections were linked with *Legionella* sp. [12], free-living amoeba (*Acanthamoeba* sp. and *Naegleria fowleri*) [13] or other protozoa, such as *Vittaforma* sp. and fungi *Ochroconis gallopava* [14]. A study carried out by Charles Rabkin (1984), found that thermophilic bacteria which are able to grow ≥50 °C can cause diseases like meningitis, endocarditis, and septicemia in the human population. Among these, many of the isolated bacteria were resistant to many antibiotics such as erythromycin, tetracycline, sulfamethoxazole, tobramycin and netilmicin [6]. Other study led by Jardine et al. 2017, showed the emergence of antibiotic resistance in isolates which were identified as opportunistic food bore pathogens such as *Arthrobacter* sp. and *Hafnia* sp. from hot springs [15]. Thus, the study on antibiotic resistance profile and their possible mechanisms would have an escalating scope. Hot springs are generally isolated ecosystems with primitive prokaryotic communities. Thus, finding the distribution of ARGs in an ecologically isolated thermophilic environment with fewer anthropogenic exposures would answer a lot of queries.

Sikkim, a northeastern state of India, with mountains as topological attributes, harbors 26 hot springs and many more yet to be discovered. Hot springs of Sikkim possess different temperatures ranging from 50-77°C with the diverse thermophilic prokaryotic community as reported in one of our earlier studies [16]. Widely belief of sacred healing capacity of the hot springs attracts tourist and pilgrims to take a thermal bath in the springs. Thus, it is important to determine the abundance of antibiotic resistance and metal resistance in a thermophilic prokaryotic community of hot springs to conserve these ecologically important reservoirs from further anthropogenic interventions. Maintaining an uncontaminated status of nature’s bathhouse for public use also ensures future health-related safety. In this study, the community structure of thermophilic bacteria in the hot springs was determined using the culture-independent method. The antibiotic resistance profile has been evaluated from the metagenome sequence analysis. Along with the ARGs, metal resistance and distribution of MRGs has also been evaluated by gene mining from metagenomic sequence library. This study will be important to understand the antibiotic and heavy metal resistance along with co-selection of resistance group among thermophilic bacteria. This may also highlight many unmarked revelations about hot springs and thermophilic bacteria residing in them.

## Materials and Methods

### Study site and Sampling

Water samples from two hot springs of Sikkim – *Reshi*, and *Yumthang* were investigated under this study. Hot spring *Reshi* is located on the bank of river Rangit in South Sikkim district whereas *Yumthang* hot spring is located on the bank of river Lachung in North Sikkim district. These hot springs are an attraction to locals and people from neighboring countries like Nepal, Bhutan, and Tibet. The bathing pond of *Yumthung* is remodeled and constructed with concrete and bricks, whereas the Reshi hot spring is in natural condition **Supplementary Fig.1a - b**. The physical parameters like temperature, pH, dissolved oxygen (DO), total dissolved solids (TDS) and conductivity were examined *in-situ* at the sampling site through Multi Water Quality Checker U-50 Series (Horiba, Japan). GPS coordinates were recorded using GPSMAP 78S (Garmin, USA) as per the guidelines of the manufacturer **Table 1**. For microbiological examination (culture-independent techniques), water samples were aseptically collected in sterile thermal flasks (Mega Slim, USA) in triplicates. Prior to analysis, samples were transferred to the laboratory and kept at 4°C and then processed immediately.

**Table.1.** Physical Parameters of four Hot Springs.

| Physical Parameters of two Hot Springs |  |  |  |  |  |  |  |
| --- | --- | --- | --- | --- | --- | --- | --- |
| Hot Spring | Coordinates | Temperature (in°C) | pH | Conductivity mS/cm | D.O. (mg/L) | D.O. (%) | TDS(g/L) |
| Reshi | 27°14.957'N<br>88°18.176'E | 47.4 | 6.57 | 0.935 | 7.07 | 104.09 | 0.598 |
| Yumthang | 27°47.575'N<br>88°42.516'E | 41 | 7.5 | 0.234 | 7.5 | 116.5 | 0.15 |

### Culture Independent technique

Shotgun metagenomics was performed for these two hot springs - *Reshi* and *Yumthang* hot springs based on two distantly located regions of South and North districts of Sikkim (as representative of the two districts).

### Metagenomic DNA extraction

Environmental DNA was extracted using DNeasy Power Water Kit (MOBIO Laboratories, USA) in accordance with manufacturer’s instruction. Quality of the DNA was checked on 0.8% agarose gel and DNA was quantified using Qubit Fluorometer (Thermofisher Scientific, USA), with a detection limit of 10-100ngμL^-1^.

### Metagenomic Data Analysis

Short gun metagenomic sequence data were generated using Illumina HiSeq 4000. Data quality was checked using FastQC and MultiQC software [17]. Microbial abundance was estimated using Metaphlan2 [18]. To generate the metagenome assembly, metaSPAdes [19] and IDBA_UD [20] were used with multiple kmer values set from 40 to 120 with increments of 20. Assemblies were scaffolded to get the best contigs. The assembly statistics were calculated using Quast [21]. The assembled contigs were annotated using BLASTn [22] against the NCBI(nt) custom metagenome database that included all nucleotide entries for organisms that belong to the taxa archaea, bacteria, fungi, and viruses. The gene prediction and annotation of the assembled contigs were carried out using PROKKA [23]. The predicted genes functional classification based on KEGG orthology was done using FMAP (Functional Mapping and Analysis Pipeline) [24] and COG (Cluster of Orthologous Groups) [25]. Diversity indices viz. Shannon, Chao-1 and Fisher alpha was calculated with PAST software [26]

### Mining ARGs and MRGs from Metagenome Sequence

Putative ARGs from the PROKKA predicted genes were identified with the ardbAnno V.1.0 [27] script available from the ARDB (Antibiotic Resistance Genes Database) consortium using the non-redundant resistance genes as a reference. Putative MRGs were identified by using BacMetScan V.1.0 [28] script available from the BacMet AntiBacterial Biocide and Metal resistance gene database. A manually curated database of genes with experimentally confirmed resistance function was used in BacMet-Scan as reference. Predicted resistant gene classification was also done based on COG (Cluster of Orthologous Groups) classification by BLASTx against the COG database [25].

## Data availability

Raw metagenomic reads were submitted to Sequence Read Archive (SRA), NCBI to obtain the Bio-Sample and Sequence Read Archive (SRA) accession numbers The accession numbers obtained are (SAMN08940114, SRP140681, and SRS3178356) for Reshi Hot Spring with sample name as RESMETAV4 and (SAMN08940367, SRP140682, and SRS3178357) for Yumthang Hot Spring with sample name as YUMMETAV4. The SRA records will be accessible with the following links for Reshi and Yumthang Hot Springs respectively https://www.ncbi.nlm.nih.gov/sra/SRP140681 and https://trace.ncbi.nlm.nih.gov/Traces/sra/?run=SRR7015754. The whole-genome Bio-sample ID’s and Genome Accession numbers are SAMN08933550/QCWL00000000 and SAMN07653191/NWUZ00000000 for LYN3 and AYN2 respectively.

### ARG and MRG analysis by Whole Genome Sequencing

The previous reported whole genome of thermophilic bacteria *Geobacillus yumthangensis* from our study [29], was used to evaluate the presence of ARGs and MRGs as a reference to prove the hypothesis. The whole-genome sequencing for another isolate, i.e., *Geobacillus toebii* LYN3 was performed using Illumina HiSeq 4000 sequencing technology with a paired-end sequencing module. The genome annotation and functional prediction were done using Rapid Annotations using Subsystems Technology RAST (V 2.0) [30].

### Statistical Analysis

Estimation of alpha diversity indices such as Shannon, Sampson, and Chao was calculated using Past software. Post hoc analysis of variance (ANOVA) and correlation analysis (Pearson and Spearman) was performed by using Xlstat and statistical analysis software - Graph Pad Prism for hypothesis testing.

## RESULTS

### Physicochemical analysis of hot spring samples

Hot springs of Sikkim are in Himalayan Geothermal Belt (HGB). Temperature profiling showed that *Reshi* hot spring was hotter (47.4 **°**C) followed by *Yumthan*g hot spring (41 **°**C) **Table 1.** The pH of hot springs *Reshi* (6.57) and *Yumthang* (7.5) was more or less neutral. The conductivity and TDS were higher 0.93 mS/cm and 0.59 (g/L) in case of *Reshi* hot spring respectively.

### Microbial diversity indices

Illumina HiSeq generated sequence data of 61898482 reads in *Reshi* hot Spring sample and 37338796 reads in *Yumthang* hot Spring sample with a read length of 150. The data was checked for base call quality distribution, and % Bases >Q20 was calculated to be equal to 98.05% and 99.26% for *Reshi* and *Yumthang* hot springs respectively. The GC content was found to be 64 and 65 for *Reshi* and *Yumthang* hot springs respectively. The estimates of alpha diversity indices revealed significant differences between Reshi and Yumthang hot springs. Shannon index for bacterial communities showed a significant variation in *Reshi* (1.09) and *Yumthang* (0.17). Similarly, the Sampson index follows the same trend with higher values for Reshi (0.5) followed by Yumthang (0.05). However, the species richness was found to be quite similar for *Reshi* (19) and *Yumthang* (18). These results indicate that *Reshi* hot spring possesses higher diversity and evenness than that of *Yumthang* hot spring as also depicted by rarefaction curve **Fig. 1**. However, overall diversity is low in both the hot springs **Table 2.** The gene prediction and annotation of the assembled contigs were carried out using PROKKA as given in **Table 3.**

**Fig. 1.**
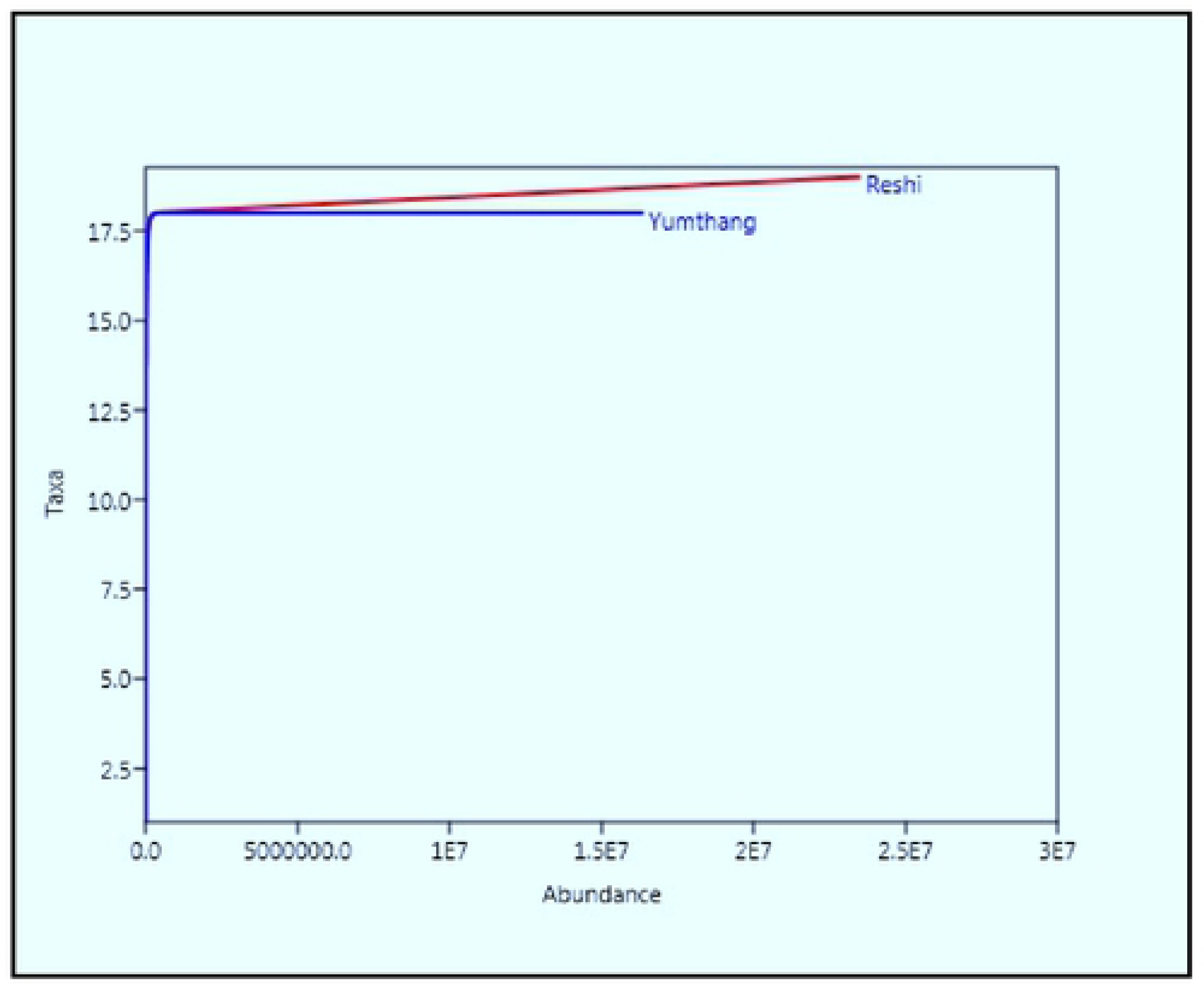
Rarefaction curve.

**Table.2.** Alpha diversity indices estimation.

| Alpha diversity indices estimation |  |  |  |  |
| --- | --- | --- | --- | --- |
|  | Shannon | Simpson | Fisher alpha | Chao-1 |
| Reshi | 1.09 | 0.57 | 1.12 | 19 |
| Yumthang | 0.17 | 0.05 | 1.08 | 18 |

**Table.3.**
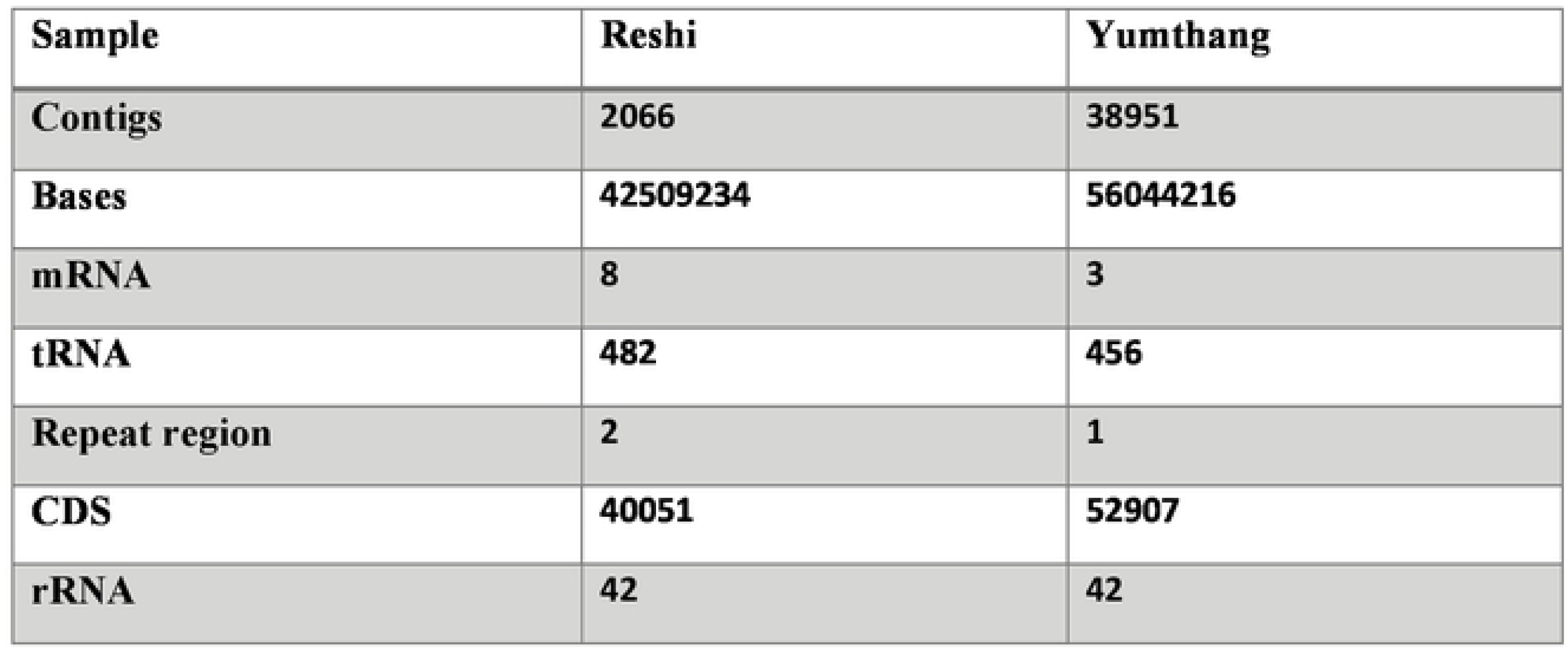
PROKKA gene prediction summary.

| Sample | Reshi | Yumthang |
| --- | --- | --- |
| Contigs | 2066 | 38951 |
| Bases | 42509234 | 56044216 |
| mRNA | 8 | 3 |
| tRNA | 482 | 456 |
| Repeat region | 2 | 1 |
| CDS | 40051 | 52907 |
| rRNA | 42 | 42 |

### Bacterial community profile

The phylum wise distribution of metagenomics data showed the dominance of *Actinobacteria* (98.1%), *Proteobacteria* (1.7%), *Firmicutes* (0.01%) and *Bacteroidetes* (0.003%) in *Yumthang* hot spring. Whereas, *Reshi* was dominated with *Proteobacteria* (75.92%), *Actinobacteria* (22.68%), *Firmicutes* (1.14%), and *Cyanobacteria* (0.03%) **Fig 2a – b.** Genus level distribution showed a distinct variation. In case of *Yumthang* hot spring the dominant genus were *Rhodococcus* (97.6%), *Escherichia* (0.73%), *Serratia* (0.49%), *Nocardiopsis* (0.47%), *Brevundimonas* (0.21%) and *Acinetobacter* (0.15%). However, in case of *Reshi* hot spring, the dominant genus was *Pseudomonas* (85.29%), *Rhodococcus* (4.28%), *Dietzia* (3.8%), *Arthobacter* (3.6%), *Staphylococcus* (1.03%) and *Paracoccus* (0.28%) **Fig 3a – b.**

**Fig. 2a.**
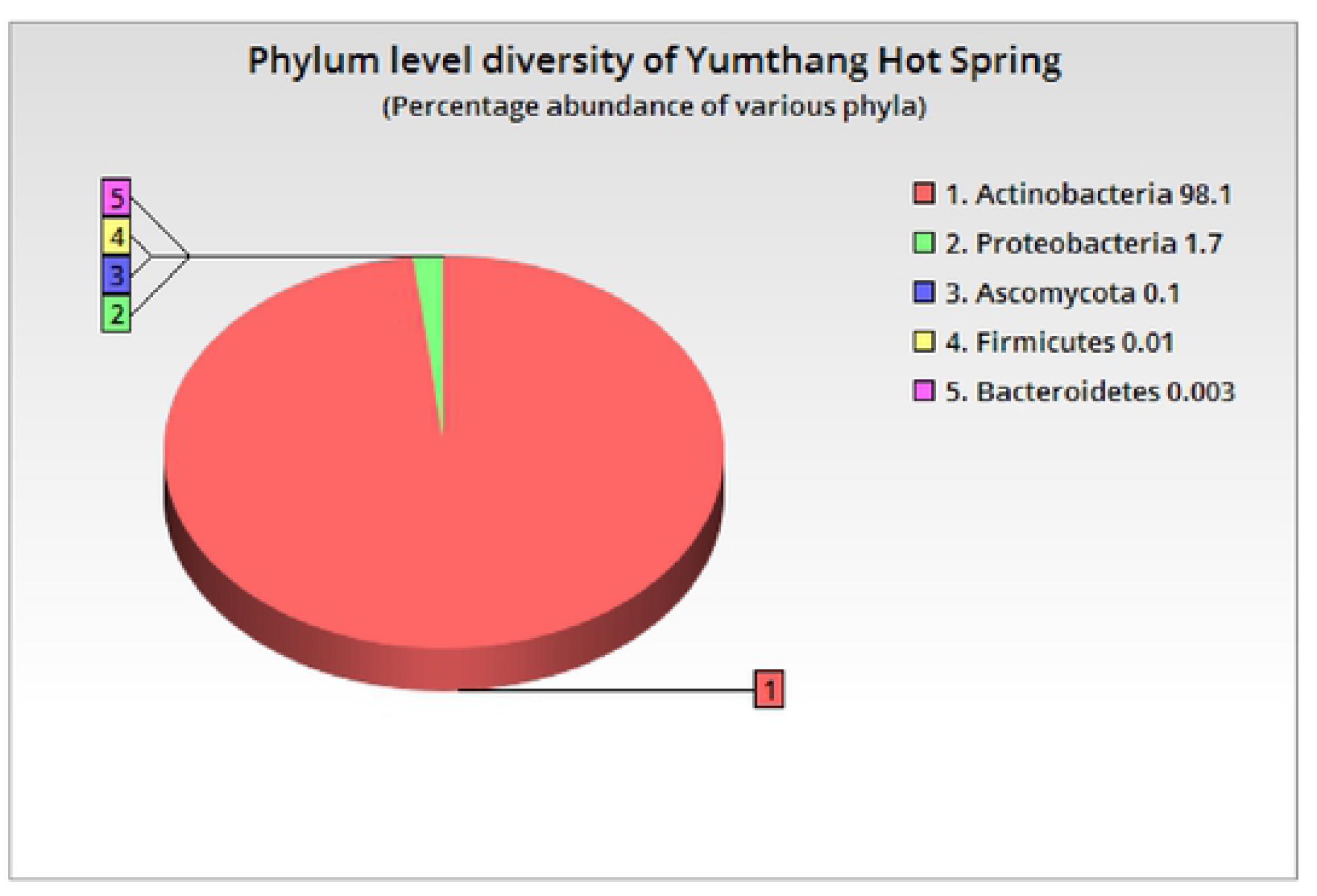
Phylum level diversity of Yumthang Hot Spring.

**Fig. 2b.**
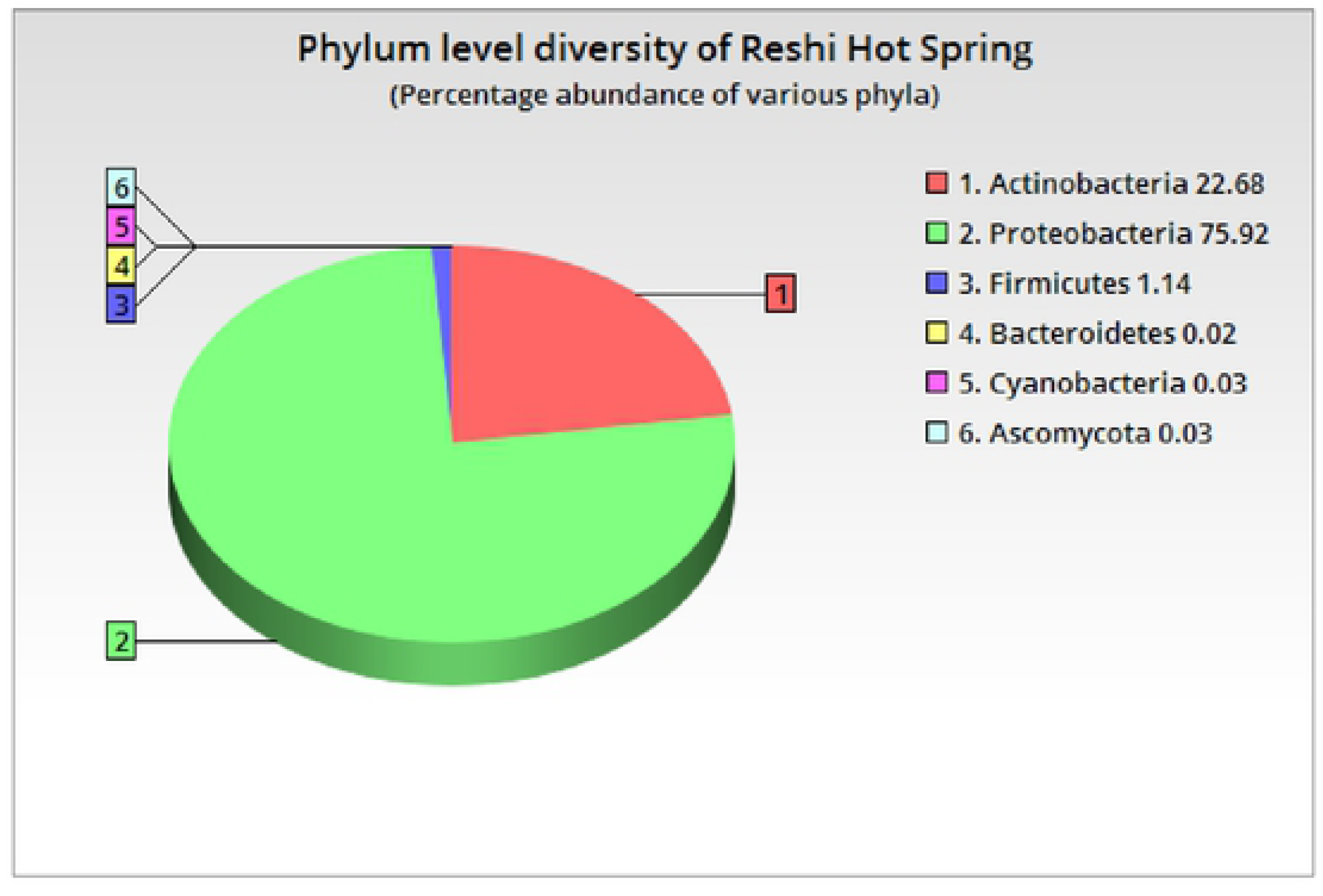
Phylum level diversity of Reshi Hot Spring.

**Fig. 3a.**
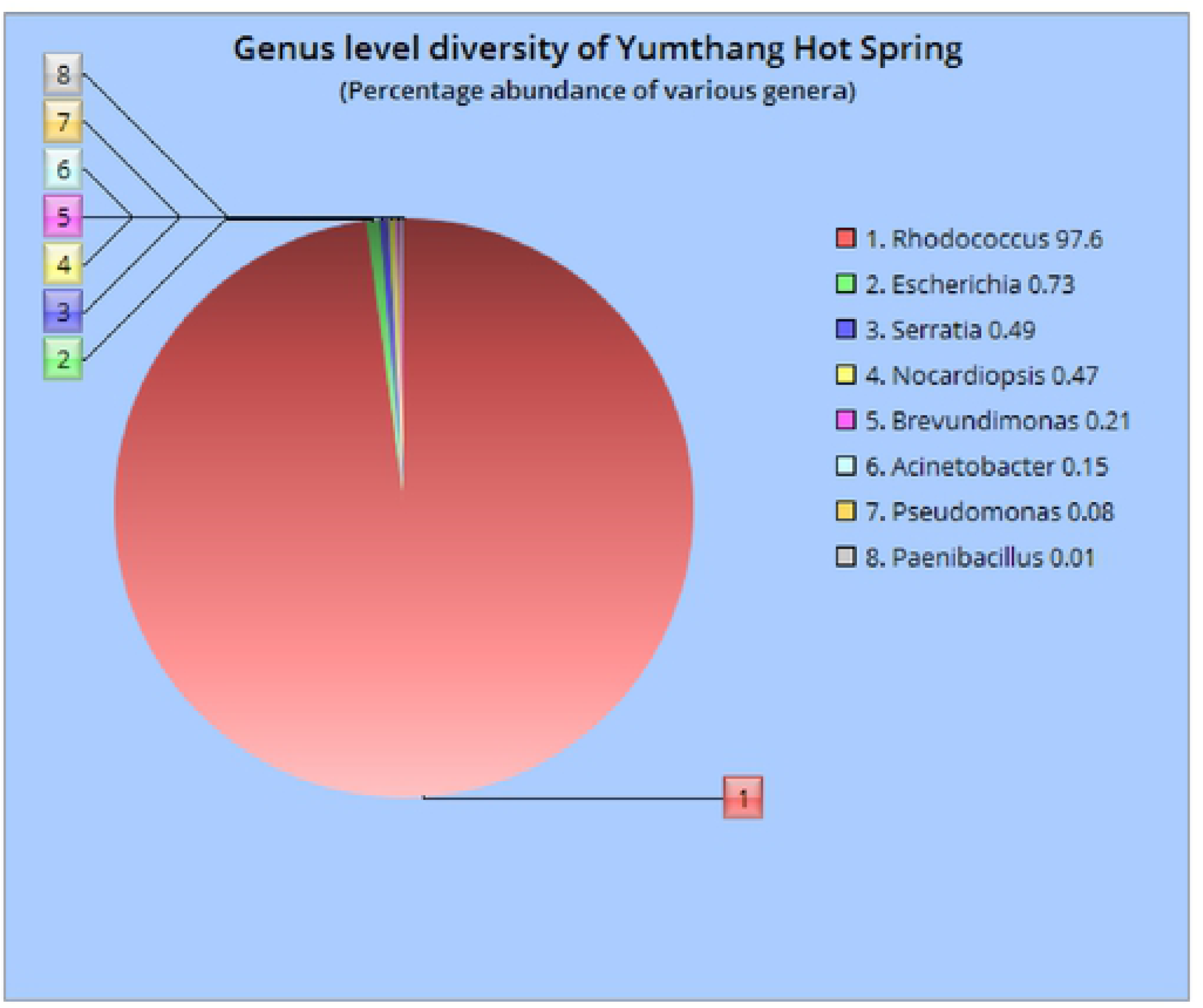
Genus level diversity of Yumthang Hot Spring.

**Fig. 3b.**
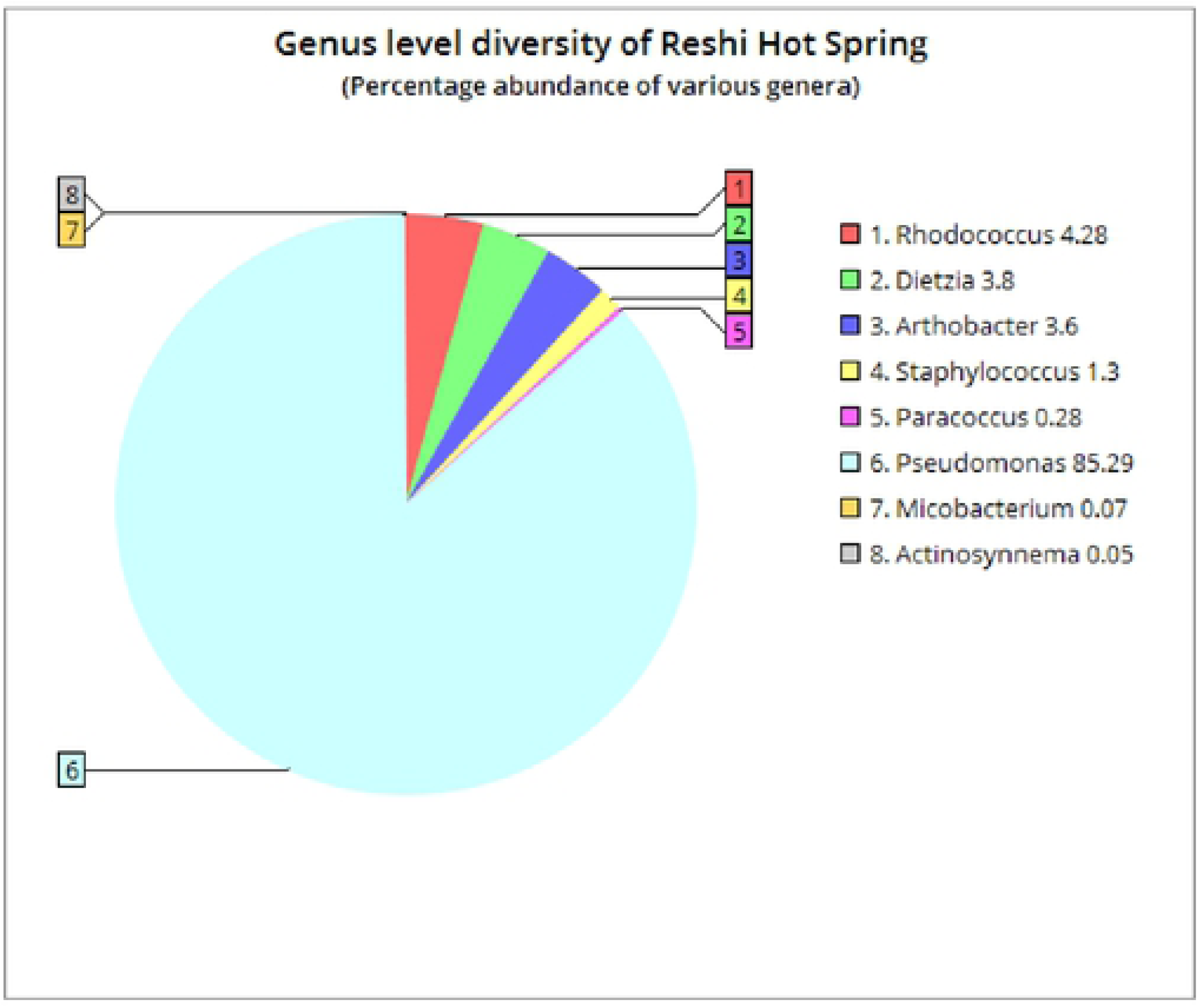
Genus level diversity of Reshi Hot Spring.

### Metagenomic studies of ARGs

Functional analysis of metagenomic ARGs data was carried out using ardbAnno V.1.0. The results have shown the presence of different antibiotics resistance genes in both the hot springs. In *Reshi* hot spring, the antibiotic resistance genes for aminoglycosides, chloramphenicol, and tetracycline were detected. The aminoglycoside gene showed the closest identity (99%) with *Acinetobacter baumanni* (genes for aminoglycoside nucleotide-transferase ANT2-1a), chloramphenicol gene (Chloramphenicol acetyltransferase) was showing (100%) identity with *Listeria monocytogenes cat*A gene. The tetracycline resistance gene (Quinolone resistance protein NorA) was showing (99%) identity with *Staphylococcus epidermidis* (*nor*A) gene. Similarly, in *Yumthang* hot spring the ARGs identified was from aminoglycosides, chloramphenicol, macrolide, β-lactams, polymyxin, and tetracycline. The aminoglycoside gene showed the closest identity (99%) with genes for aminoglycoside nucleotidal-transferase (ANT2-1a) *Acinetobacter baumanni* and aminoglycoside 6’-N-acetyltransferase (AAC(6′)-Ii) from *Serratia marcescens*. The chloramphenicol gene was showing (100%) identity with *Escherichia coli.* The tetracycline resistance gene showed identity (91%) with *Serratia marcescens* FMC 1-23-O-*tet* gene. The β-lactam gene showed identity (100%) with *E. coli* strain RM14721 and another gene from class C – β – lactamase - cephalosporin (97%) from *Serratia* sp. YD25. The macrolide gene *mac*B showed (99%) identity with *Serratia* sp. YD25 and polymyxin resistance gene showed (100%) identity with *E. coli* strain RM14721. Based on COG classification, various resistance genes were predicted corresponding to each antibiotic class as discussed above **Fig. 4.** Antibiotic classes of β-lactam showed identity with COG predicted genes such as *amp*C, *blh, Cph*A, *blab, bla, bla*Z, *hcp*A, and *pen*PC; polymyxin with COG predicted genes *arn*A; Macrolide with genes *mac*A, *mac*B and Bicyclomycin with predicted gene *bcr*.

**Fig. 4.**
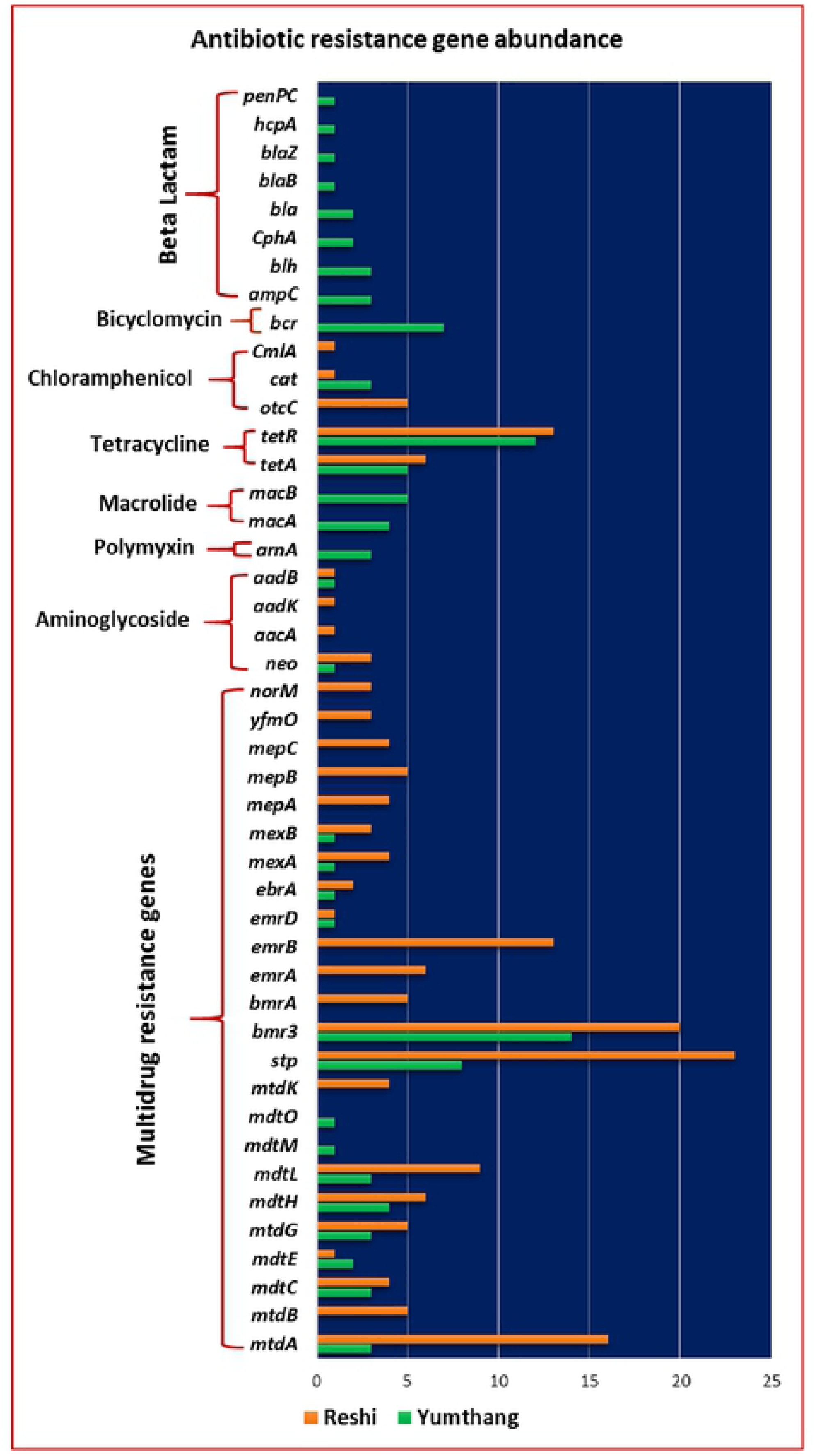
Abundance of antibiotic resistance genes. The genes were predicted from the ARDB [Antibiotic Resistance Genes Database) and resistant gene classification was also done based on COG (Cluster of Orthologous Groups.

### Metagenomic studies of MRGs

Metagenomic study of MRGs was carried out using BacMetScan V.1.0. The results showed the differential distribution of MRGs in both the hot springs. The resistant genes for cadmium, copper, zinc, mercury, and arsenic were detected. In *Yumthang* hot spring, only genes responsible for Zn tolerance were detected and genes for the resistance of Cu, Cd, Hg and As were detected from *Reshi* hot spring. Copper tolerant genes (copper resistance protein C and D) showed closest identity (100%) with *Pseudomonas fluorescens* strain TSS CopRSCD gene cluster. Cadmium tolerance gene (Cd tolerant transcriptional regulatory protein Cad) showed 99% identity with *Staphylococcus aureus* plasmid pl258 cadmium tolerance (*Cad*A) gene. Similarly, Hg tolerant gene coding for mercuric reductase showed 99% identity with *Psychrobacter* sp. strain ANT H52 (*Mer*A) gene. Arsenic tolerance genes arsenic efflux pump protein (*ars*B), arsenate reductase (*ars*C) and operon regulatory protein (*ars*R) genes showed 82-90% similarity with *Staphylococcus xylosus*. Apart from these genes efflux pump membrane transporter *BepE* was found in *Reshi* hot spring which was showing 97% identity with *Pseudomonas fluorescens* strain MFN1032 putative cation efflux protein (*mex*F) gene. In the case of Yumthang, the Zn tolerance gene for Zn uptake regulatory protein showed 100% identity with *E. coli* HKUOPY1 chromosome. Based on COG classification, various metal tolerance genes were predicted corresponding to each metal tolerance class as discussed above **Fig 5.** It was found that the metal tolerance genes were more diverse in *Reshi* hot spring than that of *Yumthang* hot spring. Metal tolerance gene classes for Cd as detected from COG predicted genes was *czz*A, *czc*B, *czc*C, *and cad*C; Hg with genes *mer*A, *mer*R, and *mer*R1; and As with genes *ars*R1, *ars*R2 and *ars*B genes from *Reshi* hot spring. However, in the case of Yumthang hot spring, only the Zn tolerance was found with the genes such as *Zur, Zup*T, *Znu*A, *Znu*B, *Znu*C, and *Znt*B.

**Fig. 5.**
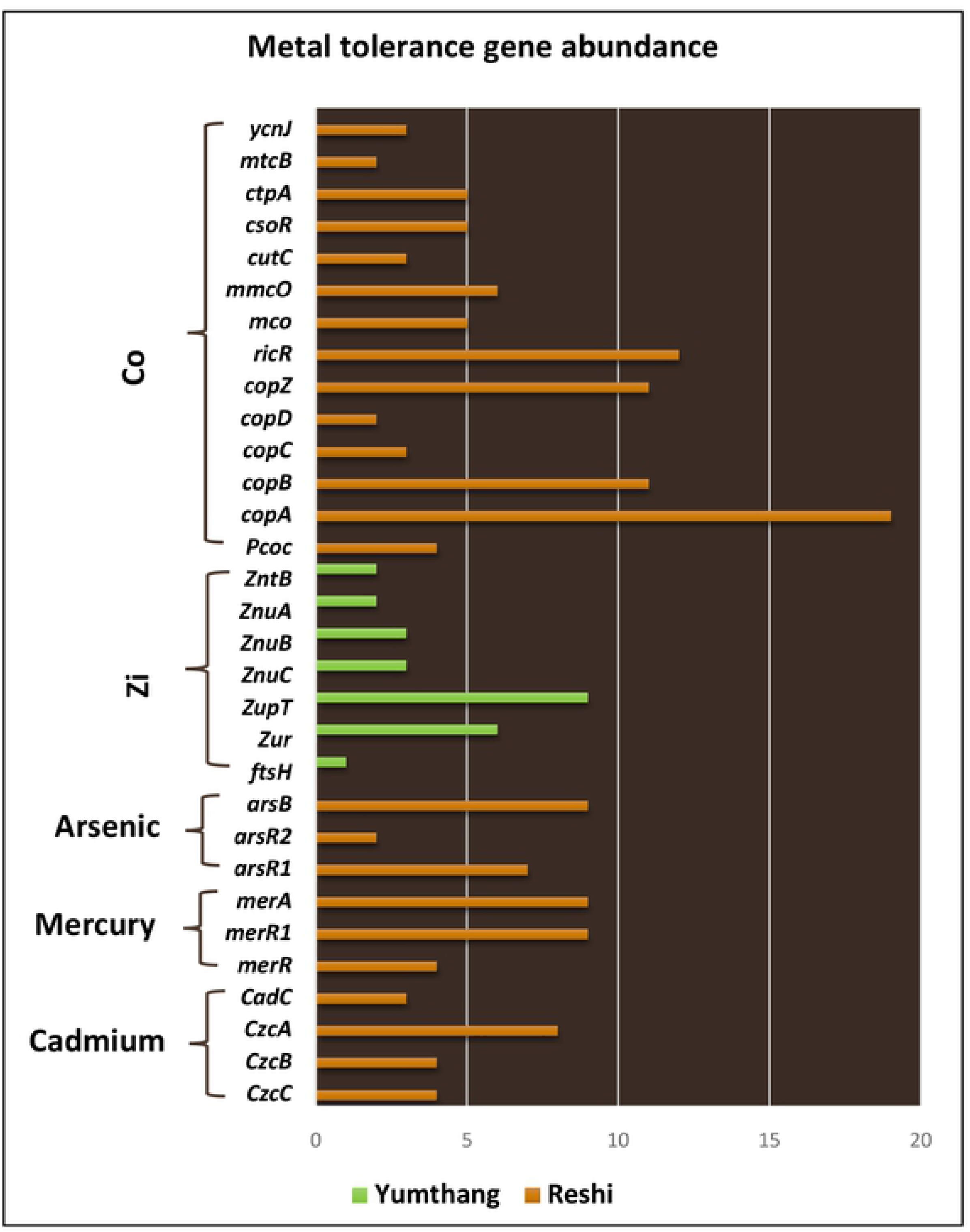
Abundance of metal resistance genes were identified from the BacMet AntiBacterial Biocide and Metal resistance gene database and resistant gene classification was also done based on COG (Cluster of Orthologous Groups) classification.

### Detection of antibiotic resistance genes and metal tolerance genes by whole-genome sequencing

The whole-genome sequence of two bacterial isolates, *Geobacillus yumthangensis* [29] and *Geobacillus toebii* LYN3 were analyzed by RAST. RAST analysis showed, both the isolate AYN2 and LYN3 were possessed putative ARGs for Fluoroquinolones, and β-lactams **Fig 6a, 6b and Supplementary Table 1.** In the case of Fluoroquinolones, four genes *par*C, *par*E, *gyr*A and *gyr*B were present which codes for DNA topoisomerase and DNA gyrase. In MRGs, genes related to copper homeostasis was found in both the isolates as copper translocating P-type ATPase or copper/silver efflux P-type ATPase. Besides, copper homeostasis, both the isolates possessed Co-Zn-Cd resistant genes. Identified genes were Co-Zn-Cd resistance protein *Czc*D; probable Co/Zn/Cd efflux system membrane fusion protein; a transcriptional regulator, *mer*R family; heavy metal resistance transcriptional regulator *hmr*R and cadmium-transporting ATPase. The strain AYN2 possessed arsenic tolerance genes for arsenical resistance operon repressor (ArsR), arsenic efflux pump protein (ArsB), arsenic reductase (ArsC), and arsenic resistance protein (ACR3). However, only ArsB, ArsC was detected in strain LYN3. Both the isolates possessed cadmium tolerance genes, cadmium transporting ATPase and cadmium efflux system accessory protein. The strain AYN2 also possessed mercury tolerance genes, Hg resistance operon regulatory protein and mercuric ion reductase, which, was found to be absent in strain LYN3.

**Fig. 6a.**
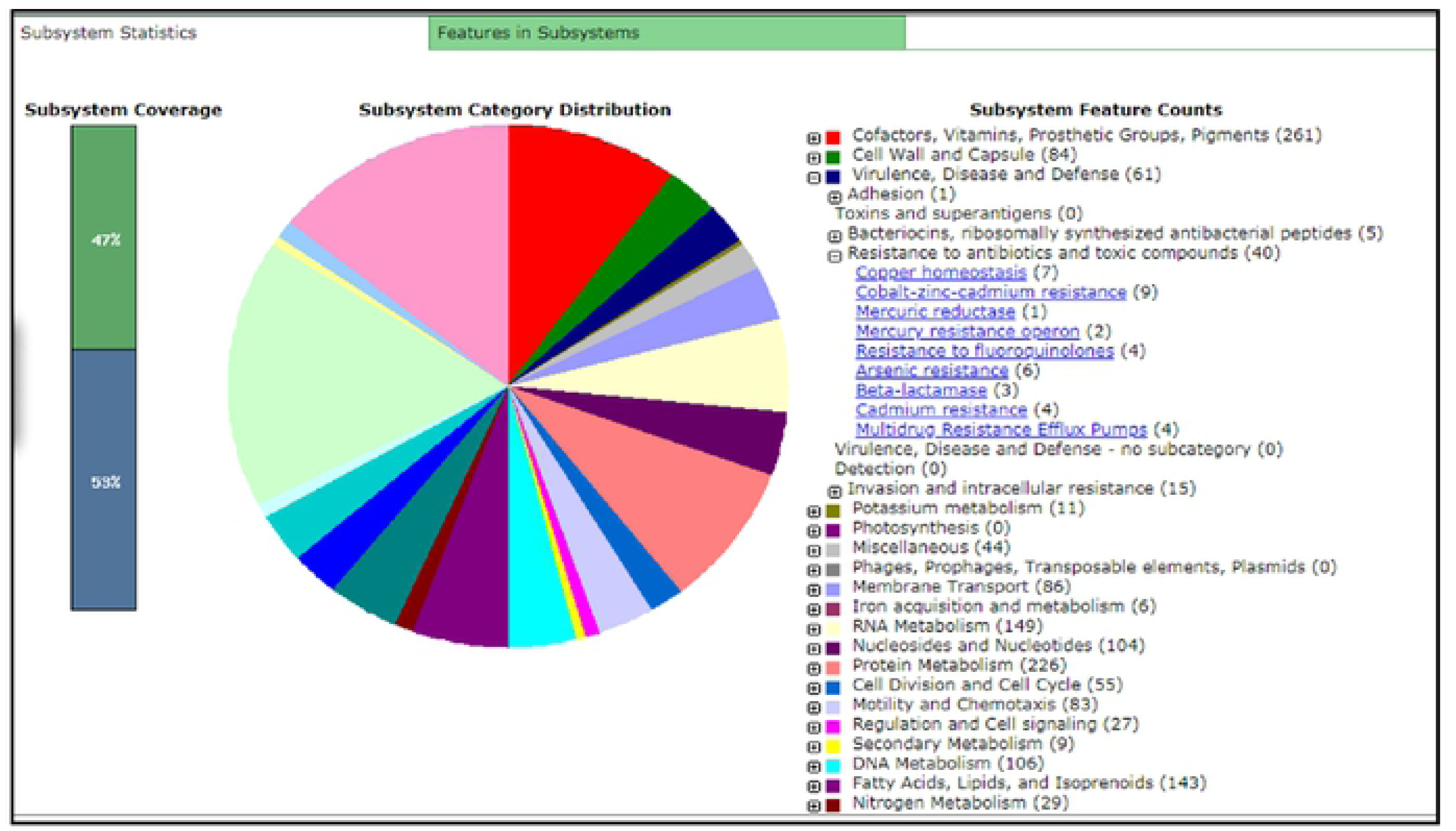
Subsystem features of Geohacillus *yumthangensis* AYN2 showing antibiotic resistance.

**Fig. 6b.**
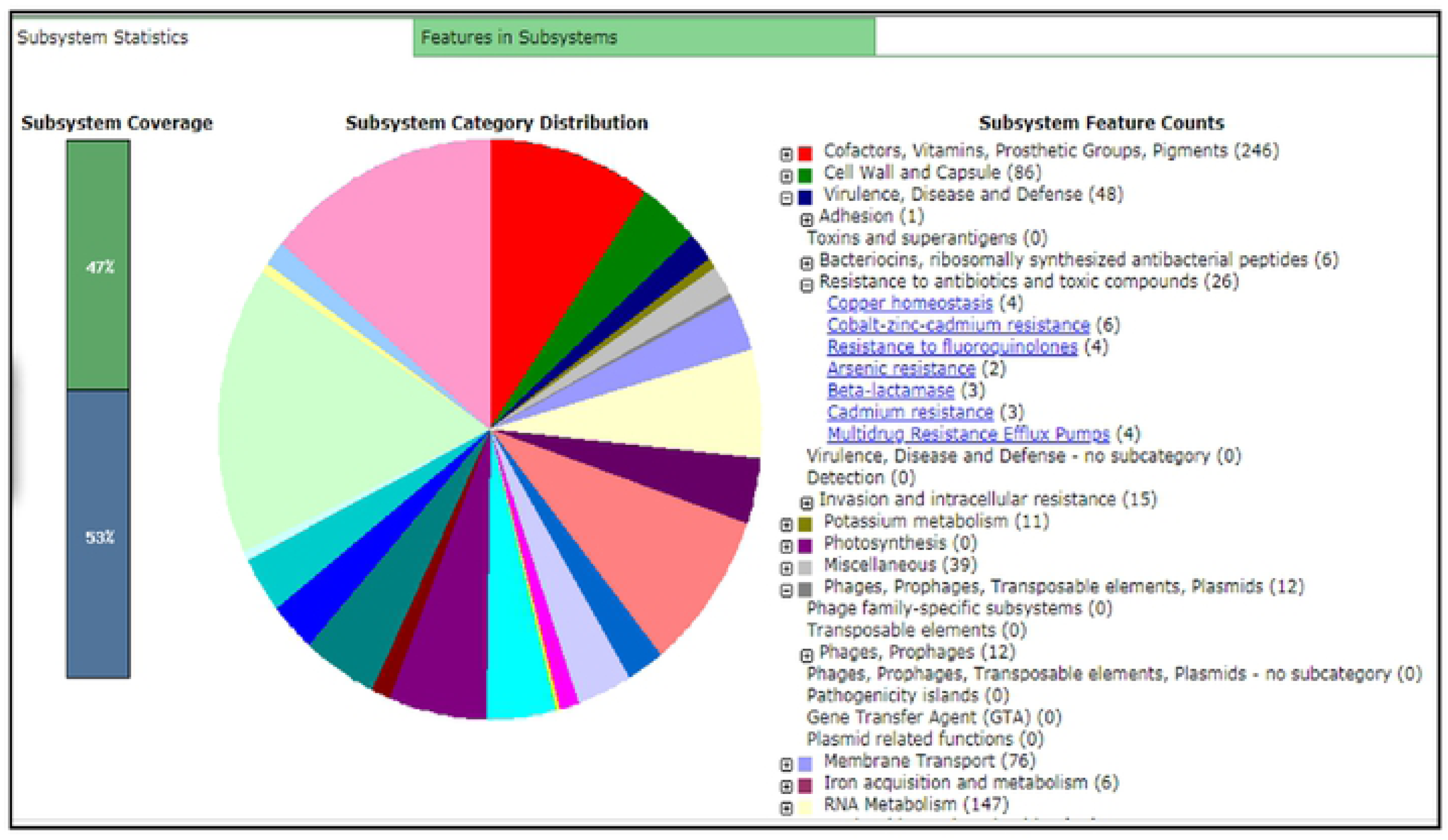
Subsystem Features of *Gieobacillus* LYN3 showing antibiotic resistance.

## Discussion

The early discovery of penicillin by Alexander Fleming and other antibiotics in the later time period has remarkably increased the life expectancies of human and improvement of human health to such an extent that in 1967 Dr. William Stewart stated that “The time has come to close the book on infectious diseases” [31]. However, exponential growth of antibiotic resistance in pathogenic and non-pathogenic bacteria in the recent era has started a wave of concern. An increasing rate of antibiotic resistance in microorganisms is critical and a threat to human health [2]. The environmental microbes which neither causes disease nor closely related to antibiotic production play a very important role in the evolutionary development of antibiotic resistance. It has been proposed that naturally the pathogens are susceptible to antibiotics or they don’t carry antibiotic-resistant genes, however they acquire it from environmental microbes. Mechanism of evolution and proliferation of ARGs in pathogens or environmental microbes is still elusive. Antibiotic-resistant bacteria and resistant genes are found in every habitat including isolated ecosystem of hundreds or thousand years [32–34]. Pathogenic and non-pathogenic bacteria belonging to the mesophilic and psychrophilic environment has been widely studied for antibiotic resistance. However, antibiotic resistance of thermophilic bacteria is yet to grasp attention. This study was designed to address the distribution of ARGs in thermophilic bacteria. The bacterial diversity and antibiotic resistance profile in two hot springs of Sikkim (*Yumthang*, and *Reshi*) have been assessed by independent techniques.

Metagenomic analysis showed the dominance of both Gram-negative (Proteobacteria) as and Gram-positive bacteria (Firmicutes). Despite being distant geographical location, bacterial community structure doesn’t show substantial variation between springs. Major phylum identified were Proteobacteria, Actinobacteria, Firmicutes, Chloroflexi, Bacteroidetes and Cyanobacteria from both the hot spring. However, the percentage of abundance varied. *Geobacillus* was the genus found which was also found to be a most dominant genus in both the hot springs via culture-dependent analysis (data not shown). The trivial difference in community composition may also reflect their difference in Physico-chemical parameters in addition to location. The phylum wise diversity in *Yumthang* hot spring and *Reshi* hot spring were in accordance with other two hot springs of Sikkim, i.e., *Polok* and *Borong* hot spring (studied earlier) [16]. The similarity in community structure can also be observed through other hot springs located across India [35–41] and China [42]. The metagenomic analysis also revealed the absence of any archaeal communities in these hot springs of Sikkim.

It was reported that Firmicutes and Actinobacteria are not the important hosts of ARGs and their numbers in microbial community increases with increase in temperature of the local environment [43]. *Geobacillus* sp. are among those group of bacteria which does not have myriad antibiotic marker genes, unlike other *Bacillaceae* members. Interestingly, whole-genome sequencing carried out in this study on two *Geobacillus* species results in detection of few putative resistance genes of fluoroquinolone class (*par*C, *par*E, *gyr*A and *gyr*B), β-lactamase (metal-dependent β-lactamase superfamily-I protein) and multidrug resistance efflux pumps (Acriflavin resistance protein and MATE family of MDR efflux pumps). However, these genes are housekeeping genes and similarity search with ARDA database doesn’t give any hit. Moreover, RAST results also indicated that they have other functions. For instance, *par*C and *par*E genes code for subunit-A and subunit-B of type-II ATP dependent DNA Topoisomerase-IV whereas, *gyr*A and *gyr*B code for subunit-A and subunit-B of DNA gyrase. DNA topoisomerase and DNA gyrase are essential genes and are involved in cell division and DNA replication. Interestingly the above discussed four genes and metal-dependent β-lactamase superfamily-I proteins along with multidrug resistance efflux pumps (Acriflavin resistance protein and MATE family of MDR efflux pumps) are present in most of the *Geobacillus* species (NCBI accession no: CM002692.1; CP002835.1; CP017694.1; CP008903.1; CP001638.1; CP017692.1 and CP016552.1). *Geobacillus yumthangensis* and *Geobacillus sp. lyn* in spite of having these genes were susceptible to at least 10 antibiotics (unpublished data) and thus it indicates that these genes have other functions in cell metabolism and can be considered as putative and inactive with respect to antibiotic resistance. Resistance genes for β-lactam antibiotic are not surprising and almost expected, as β-lactamases are involved in common bacterial functions of cell wall biosynthesis, signaling molecules, detoxification or metabolites and other functions [44].

Also, the presence of similar genes in other *Geobacillus* species and absence of most of antibiotic resistance genes suggest that *Geobacillus* species which are one of the major residents of hot springs are usually susceptible to most of the antibiotics. It was thus interesting to check if the other residents like *Anoxybacillus, Bacillus* and other thermophiles of hot springs also have these genes and absence of major antibiotic resistance genes. Research carried out by various researchers on *bacillus* indicate the similar results that these isolates are susceptible to most of the antibiotics checked. A study carried out by Coonrod et al. (1971) showed that 49 *bacillus* isolates were tested against six classes of antibiotics [45]. All were sensitive to tetracycline, kanamycin, gentamycin, and chloramphenicol. Results of this study are similar to the findings reported by Coonrod et al. (1971). Similarly Jardine et al. (2019), studied three *Bacillus* strains isolated from Jordanian hot springs against 18 antibiotics. They have also shown that the resistance was found to be less than 10% or completely sensitive. Their results also corroborate the AR profiles reported for other hot springs and confirm that lower AR levels in their studied hot springs than those reported in other environments considered to be pristine such as remote cave microbiomes, Antarctic marine waters and pristine mountain rivers [46].

Susceptible nature of these culturable bacteria may be due to many reasons such as less anthropogenic activities, high temperatures, less diversity and less competition among microbial communities. It was reported that Firmicutes and Actinobacteria are not the important hosts of ARGs and their numbers in microbial community increases with increase in temperature of the local environment [43]. *Geobacillus* sp. are among those group of bacteria which does not have myriad antibiotic marker genes, unlike other *Bacillaceae* members. Further, unavailability of any report on antibiotic-resistant *Geobacillus* sp. from the thermophilic environment along with our results suggests that naturally or generally *Geobacillus* sp. are antibiotic susceptible.

Hot springs are less intervene ecosystem from anthropogenic activities and have a high temperature. This may be the possible reason for absence of any stable antibiotics. Several authors reported that the environment with high temperature carries less antibiotic compared to normal temperature [47,48]. Thus lack of antibiotics may have been the reason for less selection pressure or competition to acquire antibiotic resistance genes by these thermophiles. These arguments are mainly based on a few factors influencing antibiotic resistance such as contact rate, transfer rate, and coexistence. Contact rates depend on the population size of the elements which communicate or transfer the gene. The abundance of only few phyla and relatively less diversity (based on very low OTUs and richness) found, suggests the low contact rate among the microbes in these hot springs. The transfer of the resistant trait is influenced by the presence of transfer elements like plasmids, chromosomal fragments or conjugative elements between bacterial cells [49]. Besides transfer element, phylogenetic closeness tends to increase HGT and stable integration [50]. However, in this study we have not detected any plasmids from the WGS of isolates and any genes which can act completely for antibiotic resistance. The genes identified which remotely can act as ARGs if they undergo mutation are also related to mesophiles and phylogenetically distant from thermophiles. Thus, the transfer rates and integration or acquisition of antibiotic resistance genes among thermophiles is very less corresponding to these hot springs. Jardine et al. 2017, showed the emergence of antibiotic resistance in isolates which were identified as opportunistic food bore pathogens such as *Arthrobacter* sp., *Kocuria* sp. and *Hafnia* sp [15]. However these three isolates have been isolated at 37°C and also literature search suggests that none of these species or genus are ever reported from hot springs, which possibly indicate that the thermophiles from hot springs are antibiotic susceptible. Exploration of antibiotic resistance genes in the hot springs of Sikkim by metagenomic analysis also suggested the limited presence of ARGs. Identified ARGs were correspondent to only Gram-negative and mesophilic bacteria (identity > 97%). Thus, it can be presumed that the genes detected through metagenomics might be intrinsic to the environment. It may also be assumed that this might possibly be the contamination of surrounding soil microflora or the human skin flora (contaminated during bathing practice). A direct correlation between human activity and antibiotic resistance has also been shown in remote Palmer sampling station in the Antarctic by Miller et al. (2009) [51].

Selection pressures like ecosystem and growth condition often determine the metabolic or physiological properties of an organism [49]. One of the selective forces we tried to find was the influence of heavy metal tolerance. Hot spring being the geological crater have a wide distribution of metallic compounds [3,17,34,39]. Besides, some of the researchers also suggested the coexistence of both ARGs and MRGs [1,52,53]. However, very less ARGs or hardly any true representative of any antibiotic resistance gene in this study decline the fact of co-existence of ARGs and MRGs in the studied thermophilic environment. In this study, we have detected metal tolerance via whole-genome sequencing of isolates AYN2 and LYN3 and these have shown the presence of MRGs for Cu, Co, Zn, Cd and As. The functional metagenomic analysis depicts the similar results with identified MRGs viz czzA, czcB, czcC, and cadC, merA, merR, merR1, arsR1, arsR2, arsB, Zur, ZupT, ZnuA, ZnuB, ZnuC, and ZntB. The hot spring water comes out of reservoirs deep down the earth through fissures surrounded by various rocks with different metal. Thus, selection pressure is a driving force in an environment where metallic elements are common. Physicochemical studies by our group showed the presence calcium, magnesium, iron and zinc metal in the hot springs of Sikkim [16]. The results didn’t show any correlation among ARGs and MRGs statistically analyzed by Pearson and Spearman correlation. Detection of MRGs in contrast to their antibiotic counterparts in this study indicates negative correlation in co-selection of these genes in such habitats. However, this is a broad assumption which needs to get validated by research in such habitats. Similarly, the absence of antibiotic resistant bacteria and ARGs from studied thermophilic bacteria indicates towards the feeble stability of ARGs in thermophilic environment. Firmicutes (*Geobacillus)* are generally devoid of ARGs, which possibly supports the hypothesis that anthropogenic activities are responsible for emergences of resistant pathogens. Moreover, these environments could possibly provide ideal conditions for the study of evolution and proliferation of antibiotic resistance in environmental microbes.

### Conclusion

An eccentric selection for baseline levels of antibiotic resistance in real-time is to identify pristine sites not yet ruined by human encroachment or “belonging to an original state or condition”. As hot springs are being considered as pristine environmental niches by various researchers as the water is not stagnant and has a constant flow in one direction. Thus the present study provides an extensive framework of bacterial diversity among the two frequently visited two hot springs of Sikkim. The estimation of diversity suggests the higher diversity in Reshi hot spring than that of Yumthang hot spring. However, the overall diversity in both the hot springs is low with moderate evenness. The major phylums found are commonly present and have been reported by researchers among such ecosystems all over the world. Antibiotic resistance is threatening mankind. Setback in the discovery of novel parvo or development of a new class of antibiotics has alarmed the scenario. Identification of ARGs in environmental bacteria and their distribution pattern has grasp attention to determine future management methods. However, the origin and evolution of the resistomes are still in ambiguity and requires much more studies. The study of ARGs and MRGs among culturable and non-culturable bacteria from hot spring of Sikkim via metagenomics showed a preferential selection of MRGs over ARGs. The absence of ARGs also declines the fact of co-selection of ARGs and MRGs in these environments. Less anthropogenic activity and human intervention of the hot springs reduces the possibility of external antibiotic pressure on the indigenous microflora. Other factors such as temperature dependence of antibiotics, contact rates or Horizontal gene transfer may affect the antibiotic susceptibility in such an ecosystem. This discrepancy between “pristine” AR levels suggests that further investigation is required in other less antibiotic exposed habitats like thermal vents, high-temperature oil fields to define baseline intrinsic levels of environmental antibiotic resistance and thus may shed further light on this subject.

## Acknowledgment

We are grateful to the Forest Department, Govt. of Sikkim for providing research permit and access to the sampling sites. We are thankful to the Department of Microbiology, Sikkim University for providing infrastructure to conduct this research.

## Declarations

### Author’s Contribution

NT designed the study, INN did the experimental works, MTS and SKD helped in the sample collection and field study, INN and SD did the analysis and prepared the manuscript, NT reviewed and edited the manuscript.

### Ethics approval and consent to participate

Not applicable

### Consent for publication

Not applicable

## References

1. Dafale NA, Semwal UP, Rajput RK, Singh GN. Selection of appropriate analytical tools to determine the potency and bioactivity of antibiotics and antibiotic resistance. J Pharm Anal. 2016;6:207–213. doi:10.1016/j.jpha.2016.05.006

2. Miller JH, Novak JT, Knocke WR, Pruden A. Survival of antibiotic resistant bacteria and horizontal gene transfer control antibiotic resistance gene content in anaerobic digesters. Front Microbiol. 2016;7:1–11. doi:10.3389/fmicb.2016.00263

3. Roca I, Akova M, Baquero F, Carlet J, Cavaleri M, Coenen S, et al. The global threat of antimicrobial resistance: Science for intervention. New Microbes New Infect. 2015;6:22–29. doi:10.1016/j.nmni.2015.02.007

4. Ventola CL. The antibiotic resistance crisis: part 1: causes and threats. P T. 2015;40:277–83. doi:Article

5. Tacconelli E, Carrara E, Savoldi A, Harbarth S, Mendelson M, Monnet DL, et al. Discovery, research, and development of new antibiotics: the WHO priority list of antibiotic-resistant bacteria and tuberculosis. Lancet Infect Dis. 2018;18:318–327. doi:10.1016/S1473-3099(17)30753-3

6. Rabkin CS, Galaid EI, Hollis DG, Weaver RE, Dees SB, Kai A, et al. Thermophilic Bacteria: New Cause of Human Disease. J Clin Microbiol. 1985;21:553–557.

7. Kurosawa H, Fujita M, Kobatake S, Kimura H, Ohshima M, Nagai A, et al. A case of Legionella pneumonia linked to a hot spring facility in Gunma Prefecture, Japan. Jpn J Infect Dis. 2010;63:78–9.

8. Iyer A, Mody K, Jha B. Biosorption of heavy metals by a marine bacterium. Mar Pollut Bull. 2005;50:340–343. doi:10.1016/j.marpolbul.2004.11.012

9. Cesare ADI, Eckert EM, Corno G. Co-selection of antibiotic and heavy metal resistance in freshwater bacteria. J Limnol. 2016;75:59–66. doi:10.4081/jlimnol.2016.1198

10. Nasermoaddeli A, Kagamimori S. Balneotherapy in medicine: A review. Environ Health Prev Med. 2005;10:171–179. doi:10.1007/BF02897707

11. Singh RN, Kaushik R, Arora DK, Saxena AK. Prevalence of opportunist pathogens in thermal prings of devotion. J Appl Sci Environ Sanit. 2013;8:195–203.

12. Sukthana Y, Lekkla A, Sutthikornchai C, Wanapongse P, Vejjajiva A, Bovornkitti S. Spa, springs and safety. Southeast Asian J Trop Med public Heal. 2005;36 Suppl 4: 10–16.

13. Fan NW, Wu CC, Chen TL, Yu WK, Chen CP, Lee SM, et al. Microsporidial keratitis in patients with hot springs exposure. J Clin Microbiol. 2012;50:414–418. doi:10.1128/JCM.05007-11

14. Yarita K, Sano A, Murata Y, Takayama A, Takahashi Y, Takahashi H, et al. Pathogenicity of Ochroconis gallopava isolated from hot springs in Japan and a review of published reports. Mycopathologia. 2007;164:135–147. doi:10.1007/s11046-007-9034-7

15. Jardine JL, Abia ALK, Mavumengwana V, Ubomba-Jaswa E. Phylogenetic analysis and antimicrobial profiles of cultured emerging opportunistic pathogens (Phyla actinobacteria and proteobacteria) identified in hot springs. Int J Environ Res Public Health. 2017;14:1–18. doi:10.3390/ijerph14091070

16. Najar IN, Sherpa MT, Das S, Das S, Thakur N. Microbial ecology of two hot springs of Sikkim: Predominate population and geochemistry. Sci Total Environ. 2018;637–638:730–745. doi:10.1016/j.scitotenv.2018.05.037

17. Ewels P, Magnusson M, Lundin S, Käller M. MultiQC: Summarize analysis results for multiple tools and samples in a single report. Bioinformatics. 2016;32:3047–3048. doi:10.1093/bioinformatics/btw354

18. Truong DT, Franzosa EA, Tickle TL, Scholz M, Weingart G, Pasolli E, et al. MetaPhlAn2 for enhanced metagenomic taxonomic profiling. Nat Methods. 2015;12:902–903. doi:10.1038/nmeth.3589

19. Nurk S, Meleshko D, Korobeynikov A, Pevzner PA. MetaSPAdes: A new versatile metagenomic assembler. Genome Res. 2017;27:824–834. doi:10.1101/gr.213959.116

20. Peng Y, Leung HCM, Yiu SM, Chin FYL. IDBA-UD: A de novo assembler for single-cell and metagenomic sequencing data with highly uneven depth. Bioinformatics. 2012;28:1420–1428. doi:10.1093/bioinformatics/bts174

21. Gurevich A, Saveliev V, Vyahhi N, Tesler G. QUAST: Quality assessment tool for genome assemblies. Bioinformatics. 2013;29:1072–1075. doi:10.1093/bioinformatics/btt086

22. Altschul SF, Gish W, Miller W, Myers EW, Lipman DJ. Basic local alignment search tool. J Mol Biol. 1990;215:403–10. doi:10.1016/S0022-2836(05)80360-2

23. Seemann T. Prokka: Rapid prokaryotic genome annotation. Bioinformatics. 2014;30:2068–2069. doi:10.1093/bioinformatics/btu153

24. Kim J, Kim MS, Koh AY, Xie Y, Zhan X. FMAP: Functional Mapping and Analysis Pipeline for metagenomics and metatranscriptomics studies. BMC Bioinformatics. 2016;17:1–8. doi:10.1186/s12859-016-1278-0

25. Tatusov RL, Fedorova ND, Jackson JD, Jacobs AR, Kiryutin B, Koonin E V, et al. The COG database: an updated version includes eukaryotes. BMC Bioinformatics. 2003;14:1–14.

26. Chao A, Chazdon RL, Colwell RK, Shen TJ. Abundance-based similarity indices and their estimation when there are unseen species in samples. Biometrics. 2006;62:361–371. doi:10.1111/j.1541-0420.2005.00489.x

27. Liu B, Pop M. ARDB - Antibiotic resistance genes database. Nucleic Acids Res. 2009;37:443–447. doi:10.1093/nar/gkn656

28. Pal C, Bengtsson-Palme J, Rensing C, Kristiansson E, Larsson DGJ. BacMet: Antibacterial biocide and metal resistance genes database. Nucleic Acids Res. 2014;42:737–743. doi:10.1093/nar/gkt1252

29. Najar IN, Sherpa MT, Das S, Verma K, Dubey VKNT. Geobacillus yumthangensis sp. nov., a thermophilic bacterium isolated from a north-east Indian hot spring. Int J Syst Evol Microbiol. 2018; 1–5. doi:10.1099/ijsem.0.003002

30. Aziz RK, Bartels D, Best AA, DeJongh M, Disz T, Edwards RA, et al. The RAST Server: rapid annotations using subsystems technology. BMC Genomics. 2008;9:75. doi:10.1186/1471-2164-9-75

31. Upshur R. Ethics and infectious disease. Bull World Health Organ. 2008;86:654. doi:http://dx.doi.org/10.1016/S0002-9394(14)70407-6

32. Clemente JC, Pehrsson EC, J.Blaser M, Sandhu K, Gao Z, Wang B, et al. The microbiome of uncontacted Amerindians. Microb Ecol. 2015;4:1–12. doi:10.1063/1.4898326

33. Goethem MW Van, Pierneef R, Bezuidt OKI, Peer Y Van De, Cowan DA, P. TM. A reservoir of ‘historical’ antibiotic resistance genes in remote pristine Antarctic soils. Microbiome. 2018;6:1–12. doi:10.1103/PhysRevD.65.065015

34. Pawlowski AC, Wang W, Koteva K, Barton HA, McArthur AG, Wright GD. A diverse intrinsic antibiotic resistome from a cave bacterium. Nat Commun. 2016;7:1–10. doi:10.1038/ncomms13803

35. Badhai J, Ghosh TS, Das SK. Taxonomic and functional characteristics of microbial communities and their correlation with physicochemical properties of four geothermal springs in Odisha, India. Front Microbiol. 2015;6:1–15. doi:10.3389/fmicb.2015.01166

36. Ghelani A, Patel R, Mangrola A, Dudhagara P. Cultivation-independent comprehensive survey of bacterial diversity in Tulsi Shyam Hot Springs, India. Genomics Data. 2015;4:54–56. doi:10.1016/j.gdata.2015.03.003

37. Mangrola A V., Dudhagara P, Koringa P, Joshi CG, Patel RK. Shotgun metagenomic sequencing based microbial diversity assessment of Lasundra hot spring, India. Genomics Data. 2015;4:73–75. doi:10.1016/j.gdata.2015.03.005

38. Mehetre GT, Paranjpe A, Dastager SG, Dharne MS. Investigation of Microbial Diversity in Geothermal Hot Springs in Unkeshwar, India, Based on 16S rRNA Amplicon Metagenome Sequencing. Genome Announc. 2016;4:4–5. doi:10.1128/genomeA.01766-15.Copyright

39. Panda AK, Bisht SS, De Mandal S, Kumar NS. Bacterial and archeal community composition in hot springs from Indo-Burma region, North-east India. AMB Express. 2016;6:111. doi:10.1186/s13568-016-0284-y

40. Sangwan N, Lambert C, Sharma A, Gupta V, Khurana P, Khurana JP, et al. Arsenic rich Himalayan hot spring metagenomics reveal genetically novel predator – prey genotypes. Environ Microbiol Rep. 2015;7:812–823. doi:10.1111/1758-2229.12297

41. Saxena R, Dhakan DB, Mittal P, Waiker P. Metagenomic Analysis of Hot Springs in Central India Reveals Hydrocarbon Degrading Thermophiles and Pathways Essential for Survival in Extreme Environments. Front Microbiol. 2017;7:1–17. doi:10.3389/fmicb.2016.02123

42. Hedlund BP, Cole JK, Williams AJ, Hou W, Zhou E, Li W, et al. A review of the microbiology of the Rehai geothermal field in Tengchong, Yunnan Province, China. Geosci Front. 2012;3:273–288. doi:10.1016/j.gsf.2011.12.006

43. Macfadden DR, Mcgough SF, Fisman D, Brownstein JS, Group E, Program HI. Antibiotic Resistance Increases with Local Temperature. Nat Clim Chang. 2018;8:510–514. doi:10.1038/s41558-018-0161-6.Antibiotic

44. Martínez JL. Natural antibiotic resistance and contamination by antibiotic resistance determinants: The two ages in the evolution of resistance to antimicrobials. Front Microbiol. 2012;3:2010–2012. doi:10.3389/fmicb.2012.00001

45. Coonrod JD, Leadley PJ, Eickhoff TC. Antibiotic Susceptibility of. J Infect Dis. 1971;123:102–105. doi:10.1159/000007304

46. Jardine J, Mavumengwana V, Ubomba-Jaswa E. Antibiotic resistance and heavy metal tolerance in cultured bacteria from hot springs as indicators of environmental intrinsic resistance and tolerance levels. Environ Pollut. 2019;249:696–702. doi:10.1016/j.envpol.2019.03.059

47. Diehl DL, Lapara TM. Effect of Temperature on the Fate of Genes Encoding Tetracycline Resistance and the Integrase of Class 1 Integrons within Anaerobic and Aerobic Digesters Treating Municipal Wastewater Solids. Environ Sci Technol. 2010;44:9128–9133.

48. Sun W, Qian X, Gu J, Wang X-J, Duan M-L. Mechanism and Effect of Temperature on Variations in Antibiotic Resistance Genes during Anaerobic Digestion of Dairy Manure. Sci Rep. 2016;6:30237. doi:10.1038/srep30237

49. Martinezi JL, Baquero F. Mutation Frequencies and Antibiotic Resistance. Antimicrob Agents Chemother. 2000;44:1771–1777. doi:10.1128/AAC.44.7.1771-1777.2000.Updated

50. Smillie CS, Smith MB, Friedman J, Cordero OX, David LA, Alm EJ. Ecology drives a global network of gene exchange connecting the human microbiome. Nature. 2011;480:241–244. doi:10.1038/nature10571

51. Miller R V., Gammon K, Day MJ. Antibiotic resistance among bacteria isolated from seawater and penguin fecal samples collected near Palmer Station, AntarcticaThis article is one of a selection of papers in the Special Issue on Polar and Alpine Microbiology. Can J Microbiol. 2009;55:37–45. doi:10.1139/w08-119

52. Becerra-castro C, Machado RA, Vaz-moreira I, Manaia CM. Science of the Total Environment Assessment of copper and zinc salts as selectors of antibiotic resistance in Gram-negative bacteria. Sci Total Environ. 2015;530–531:367–372. doi:10.1016/j.scitotenv.2015.05.102

53. Hu HW, Wang JT, Li J, Li JJ, Ma YB, Chen D, et al. Field-based evidence for copper contamination induced changes of antibiotic resistance in agricultural soils. Environ Microbiol. 2016;18:3896–3909. doi:10.1111/1462-2920.13370

